# Barley shoot biomass responds strongly to N:P stoichiometry and intraspecific competition, whereas roots only alter their foraging

**DOI:** 10.1101/2020.01.20.912352

**Authors:** Amit Kumar, Richard van Duijnen, Benjamin M. Delory, Rüdiger Reichel, Nicolas Brüggemann, Vicky M. Temperton

## Abstract

**Background and Aims:** Plants respond to various environmental stimuli, and root systems are highly responsive to the availability and distribution of nutrients in the soil. Root system responses to the limitation of either nitrogen (N) or phosphorus (P) are well documented, but how the early root system responds to (co-) limitation of one (N or P) or both (N and P) in a stoichiometric framework is not well known despite its relevance in agriculture. In addition, how plant-plant competition (here intra-specific) alters plant responses to N:P stoichiometry is understudied. Therefore, we aimed to investigate the effects of N:P stoichiometry and competition on root system responses and overall plant performance.

**Methods:** Plants (*Hordeum vulgare* L.) were grown in rhizoboxes for 24 days in the presence or absence of competition (three vs. one plant per rhizobox), and fertilized with different combinations of N:P (low N+low P, low N+high P, high N+low P, and high N+high P).

**Key Results:** Shoot biomass was highest when both N and P were provided in high amounts. In competition, shoot biomass decreased on average by 22%. Interestingly, N:P stoichiometry and competition had no clear effect on root biomass. However, we found distinct root responses in relation to biomass allocation across depths. Specific root length depended on the identity of limiting nutrient (N or P) and presence/absence of competition. Plants rooted deeper when N was the most limiting compared to shallower rooting when P was the most limiting nutrient.

**Conclusions:** Overall, our study sheds light on the early plant responses to plant-plant competition and stoichiometric availability of two macronutrients most limiting plant performance. With low N and P availability during early growth, higher investments in root system development can significantly trade off with aboveground productivity, and strong intra-specific competition can further strengthen such effects.

## INTRODUCTION

Nutrient foraging capacity of roots determines plant performance under both heterogeneous soil nutrient availability and belowground competition with neighbors (Stibbe and Märländer, 2002; Soleymani et al. 2011; Bennett et al. 2016; Reiss and Drinkwater, 2018). Given that nutrient foraging by roots is an active process (Zhang et al. 2019), it is very likely that plant biomass allocation and root system responses will be driven by the nutrient which is limiting plant growth the most (Poorter et al. 2012). It has previously been shown for many crops how eco-physiological (Gastal and Lemaire, 2002), morphological (Fransen et al. 1998), architectural (Williamson et al. 2001; Lopez-Bucio et al. 2003; Postma and Lynch, 2012; Lynch, 2013; Guo and York, 2019), and anatomical (Wahl et al. 2001; Postma and Lynch, 2011) root traits respond to nitrogen (N) and phosphorus (P) availability in soil. For instance, Wang et al. (2015) showed contrasting root morphological and physiological trait responses of canola, barley, and potato in relation to low P availability. In order to increase P uptake, canola exuded more citric acid and developed longer roots, barley increased exudation of malic acid and reduced its root surface area and total root length, whereas potato reduced the exudation of organic acids but increased the number of root tips. Overall, it is clear that root systems respond in a species-specific way to nutrient stimuli by modifying their size and architecture (Kembel et al. 2008; Wang et al. 2015; McNickle et al. 2016). However, to what extent stoichiometric N:P availability in soil affects root systems, and how the observed effect depends on the strength of intraspecific competition has been rarely tested.

For optimal plant physiology, the elemental N and P ratio in plant biomass should be relatively stable (Güsewell, 2004). Nitrogen is an integral part of most of the enzymatic machinery, and higher N than P demand in cell metabolism indicates that N limitation can severely affect plant growth and consequently biomass production. Hence, it becomes important to understand the root foraging responses to differential availability of both N and P during early plant establishment. Differences in mobility between N and P affect their availability to plants, and root responses are likely to be specific to nutrient distribution in soil. For example, P (as orthophosphate) is highly immobile in the soil and accumulates in the topsoil strata via plant residue and fertilizer inputs. Therefore, wide dispersion of lateral roots, enhanced adventitious rooting, and shallower root growth angles are among the key root responses that are associated with enhanced topsoil foraging for P (Lynch and Brown, 2001; Lynch, 2011). In contrast, N (as nitrate) is relatively mobile in the soil compared to P and moves down the soil strata with irrigation and precipitation events. Fewer crown roots in maize, for example, can potentially improve N acquisition by exploring deep soil strata, a key root system response (Saengwilai et al. 2014; Guo and York, 2019). Therefore, the coordinated uptake and utilization of both N and P are essential in relation to optimal plant growth. However, very little is known about how plants adjust their biomass allocation and root growth responses to soil N:P stoichiometry. It is further not clear how co-limitation of both N and P will direct the plant’s response for their uptake.

Root responses not only depend on soil nutrient availability but also on the presence of neighbors (whether of the same or different species) through root-root competition for available nutrients (Cahill et al. 2010; Faget et al., 2013; McNickle and Brown, 2014, Weidlich et al. 2018). This is particularly true in mono-cropping systems where there is strong intraspecific competition for soil nutrients, mainly because neighbors share the same life-history strategies and have similar resource demands. Intense competition results in a direct negative effect on plant growth and ultimately on yield. Bennett et al. (2016) have shown interactive effects of nutrients with or without inter- and intraspecific competition on plant biomass allocation and root system responses for grasses, legumes, and forbs. Further, Hecht et al. (2016) showed for barley that roots respond to greater intraspecific competition (via manipulating sowing density) by increasing root length density and specific root length through increased fine root production. Later, Hecht et al. (2019) showed that the greater root length density under intraspecific competition was attributed to greater main root numbers. Moreover, root responses to the intraspecific competition may also include root segregation and aggregation to maximize the acquisition of nutrients (Cahill et al. 2010; Weidlich et al. 2018; Zhang et al. 2019).

Regardless of understanding how the availability of either N or P and belowground competition affects plant growth (Thuynsma et al. 2016; Sun et al. 2016), it is unclear how plants integrate the responses to differential nutrient availability and the presence or absence of intraspecific competition during early growth stages. The aim of this study was twofold: (1) investigating how N:P stoichiometry in the soil solution affects plant performance and root system responses of barley (*Hordeum vulgare* L.); and (2) determining if intraspecific competition interacts with N:P stoichiometry in shaping plant performance.

We hypothesized that:

1. From the nutrient stoichiometry perspective, N is more limiting than P for plant growth and low availability of N has stronger effects than that of P on plant performance (both below- and aboveground).
2. The intraspecific competition will lead to strong nutrient depletion, resulting in overall biomass reduction per plant.
3. Root distribution and foraging strategy will be affected by N:P stoichiometry, with plants rooting deeper when N is limiting and shallower when P is limiting, and the strength of the response will be modulated by intraspecific competition.

## MATERIALS AND METHODS

### Experimental setup

The experiment was conducted in the greenhouse of the Leuphana University Lüneburg (Lüneburg, Germany, 53°14’23.8”N 10°24’45.5”E) from August 18^th^ 2017 to September 11^th^ 2017 for a total of 24 days. The average day/night temperature and relative humidity were 22.3/15.3°C and 60/73%, respectively. Briefly, a homogenous soil mixture was prepared using sand, loess soil (nutrient-poor, collected from a lignite mine near Jackerath, Germany), and peat potting soil (Nullerde, Einheitserde Werkverband e.V., Germany) in 8:2:1 ratio, respectively. Rhizoboxes (Height: 58 cm × Width: 26.6 cm × Thickness: 2 cm; volume: 3 L) were filled with ∼ 5 kg of soil mixture. Pre-germinated (pre-germination time: 24 h on a wet tissue paper) barley (*Hordeum vulgare* L. cv. Barke, Saatzucht Breun, Germany) seedlings were transplanted in rhizoboxes as shown in Fig.1. Each rhizobox received 1 seedling for absence and 3 seedlings (7.5 cm apart from each other) for the presence of intraspecific competition (hereafter competition). Rhizoboxes were placed in containers at a 45° angle and each container contained five rhizoboxes. In each container, the front rhizobox was covered with a black plastic plate and the last rhizobox was covered with a white polystyrene plate to maintain similar light and temperature conditions, respectively. Rhizobox position was randomly changed every fourth day.

**Fig. 1:**
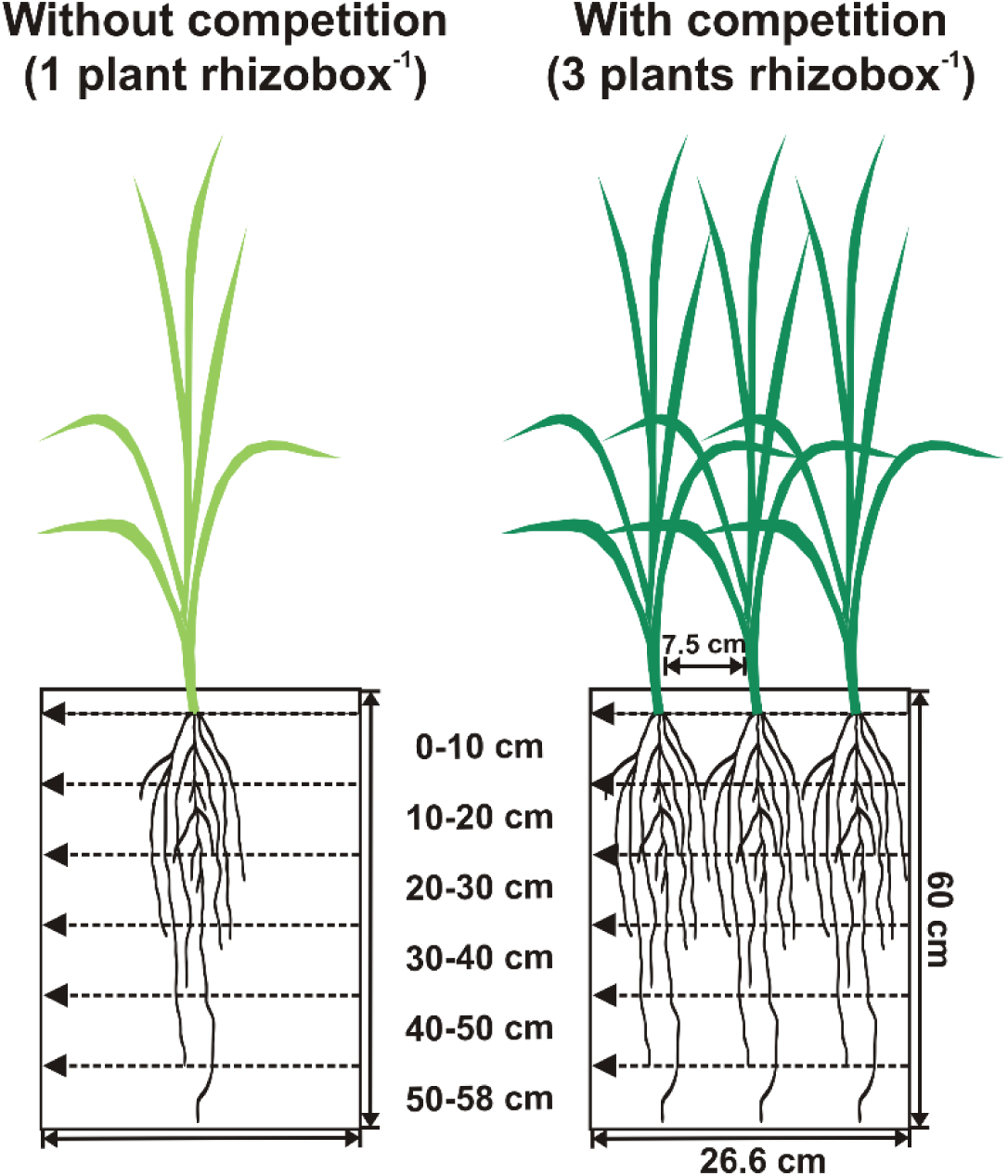
Schematic diagram representing barley grown with or without competition in rhizoboxes and showing the rooting depths sampled to assess differential root foraging responses to four N:P stoichiometry levels: low N+low P, low N+high P, high N+low P, and high N+high P.

The experiment was designed using a full factorial design to test how N:P stoichiometry (four levels: low N+low P, low N+high P, high N+low P, high N+high P) (based on pre-test showing that shoot growth was limited by N only above a ‘threshold’ of low P availability) and intraspecific competition (two levels: absence or presence of competitors) affect biomass production and allocation, soil exploration by roots, and N:P uptake of barley. In total, 8 treatment combinations were tested (4 levels of N:P stoichiometry × 2 levels of intraspecific competition) and each treatment was replicated five times resulting in a total of 40 experimental units (rhizoboxes). Rhizoboxes were provided with 800 mL of half Hoagland concentration per rhizobox before transplanting. The composition of the Hoagland solution was adjusted for each N:P stoichiometry level (low/high N, low/high P) [**Supplementary Information table 1**]. To maintain the osmotic potential, we used K_2_SO_4_ and CaCl_2_.2H_2_O as a replacement for KH_2_PO_4_, Ca(NO_3_)_2_.4H_2_O and KNO_3_ as mentioned in table 1. Rhizoboxes were left to drain for 24h and subsequently weighed. Every two days, a volume of deionized water equivalent to the evaporative loss was added to each rhizobox in order to maintain a constant weight.

### Harvest and measurements

At harvest, shoots were cut at the base and oven-dried at 80 °C until a constant mass was reached. Afterward, we carefully removed the front window of each rhizobox and divided the soil into six 10-cm layers (0-10, 10-20, 20-30, 30-40, 40-50, 50-58 cm). For each soil layer, roots were washed with tap water and stored at -20 °C until further measurements. We followed the protocol of Delory et al. (2017) for root trait measurements. Briefly, material adhering to roots was removed with brush and tweezers. In order to improve fine root detection during image analysis, clean roots were stained with a 1.7 mM neutral red solution for ∼24 h. Excess stain was removed by continuously rinsing roots with distilled water, and big root segments were cut into small pieces to avoid root overlaps during scanning. Stained roots were spread in a thin layer of distilled water in a transparent tray and scanned at 600 dpi using a commercial scanner (Epson Perfection V800 Photo, Epson, Japan). Scanned images were then analyzed with an image analysis software (WinRhizo, Regent Instruments, Quebec, Canada) using a global thresholding method. Interactive modifications to grey level pixel classification were made to improve root detection and root length estimation (Delory et al. 2017). Afterward, roots were dried at 60 °C until a constant mass was reached. Root mass fraction (RMF) was calculated as the ratio of root biomass to the total plant biomass, and specific root length (SRL) was calculated as root length per unit of root biomass.

All shoot material was ground with a ball mill (MM 400, Retsch, Germany), and measured for total C and N with an elemental analyzer (Vario EL, Elementar, Germany). For shoot P concentration, 70 mg ground samples were spiked with 2 mL HNO_3_ (65%) and 1 mL H_2_O_2_ (30%) before microwave extraction, using a MARS 5 microwave system (CEM GmbH, Germany) at 800W (80%) power, a linear temperature gradient from RT to 160°C in 20 min, holding the end temperature for 15 min. Afterward, each sample was filled up to 14 mL with ultrapure water. For P concentration determination, two aliquots of the obtained solution were diluted 1:20 with ultrapure water and analyzed. The relative standard deviation between the two repetitions was ± 10%. Total P was measured with inductively coupled plasma optical emission spectrometry (iCAP™ 7600 ICP-OES Analysator, Thermo Scientific, Germany).

### Vertical root distribution

The vertical root distribution in each rhizobox was modeled using the following asymptotic equation (Gale and Grigal, 1987; Jackson et al. 1996; Oram et al. 2018):

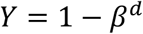

Where *Y* is the cumulative proportion [0,1] of the total root biomass located above depth *d* (in this case 0 – 58 cm), and *β* is a fitted model parameter used as a simple numerical index of vertical root distribution (Schnepf et al. 2019). Lower *β* values correspond to higher root mass allocation to surface layers, whereas higher values correspond to higher root mass allocation to deeper soil strata (Fig. 2).

**Fig. 2:**
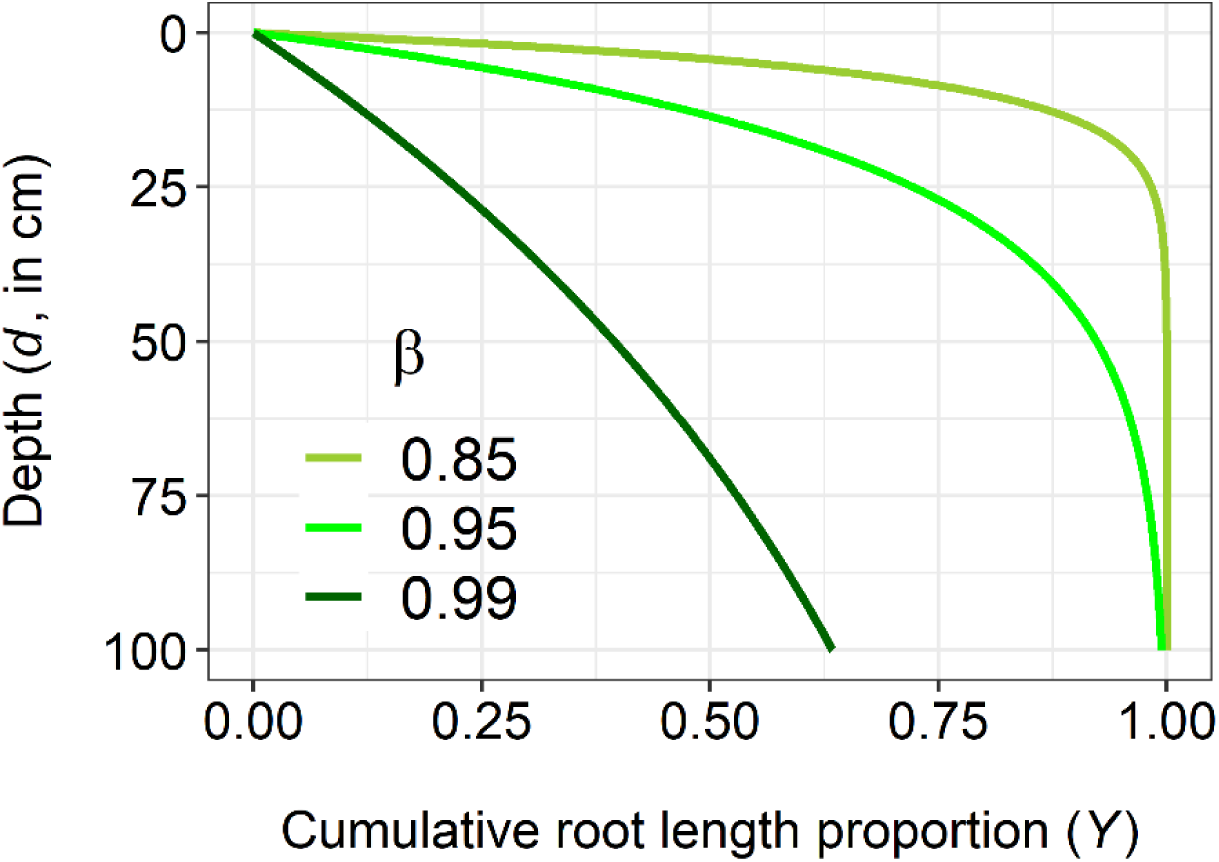
Cumulative root biomass distribution as a function of soil depth as per Gale and Grigal (1987). Higher β values imply that a greater proportion of root biomass is located in deeper soil layers, whereas lower β values imply that a greater proportion of root biomass is located in shallower soil layers.

### Statistical analyses

All statistical analyses were performed in R 3.5.0 (R Core Team, 2018) and graphs were prepared with ‘ggplot2’ library (Wickham, 2016) and R-base. The presence of potential outliers was determined graphically using the *dotchart* function. Presented in graphs are mean values of 5 replicates (4 replicates for specific root length except for LN-HP where n = 5) ± standard error (SE). Two-way ANOVA models were used to test if N:P stoichiometry, intraspecific competition, and their interaction affected shoot and root biomass, vertical root distribution (β), specific root length, and shoot N and P concentrations. Residual plots were used to check for any patterns in our data. Pairwise comparisons were performed on estimated marginal means computed by *lsmeans* using Tukey contrasts (lsmeans; Lenth, 2016). In case there was no interaction between N:P stoichiometry and competition, we show only N:P stoichiometry effects (for shoot biomass and shoot P concentration). The linear relationship between shoot N concentration and specific root length was analyzed using standard major axis (SMA) regression using the *smatr* package (Warton et al. 2012). SMA regression examines the relationship between two variables that are both measured with errors (Warton et al. 2012).

## RESULTS

### Shoot biomass

Both N:P stoichiometry (F_3,32_ = 53.08, *P* < 0.001) and competition (F_1,32_ = 52.07, *P* < 0.001) had a significant effect on shoot biomass production. The effect of N:P stoichiometry did not depend on the level of intraspecific competition (F_3,32_ = 0.48, *P* = 0.69). Looking at the effect of N:P stoichiometry, shoot biomass increased in the following order: LN-LP < LN-HP < HN-LP < HN-HP. Compared to LN-LP, shoot biomass was on average 12%, 32%, and 58% greater under LN-HP, HN-LP, and HN-HP, respectively (Fig. 3A). For plants grown in the presence of competitors, shoot biomass was on average 22% lower than plants grown in the absence of competition.

**Fig. 3:**
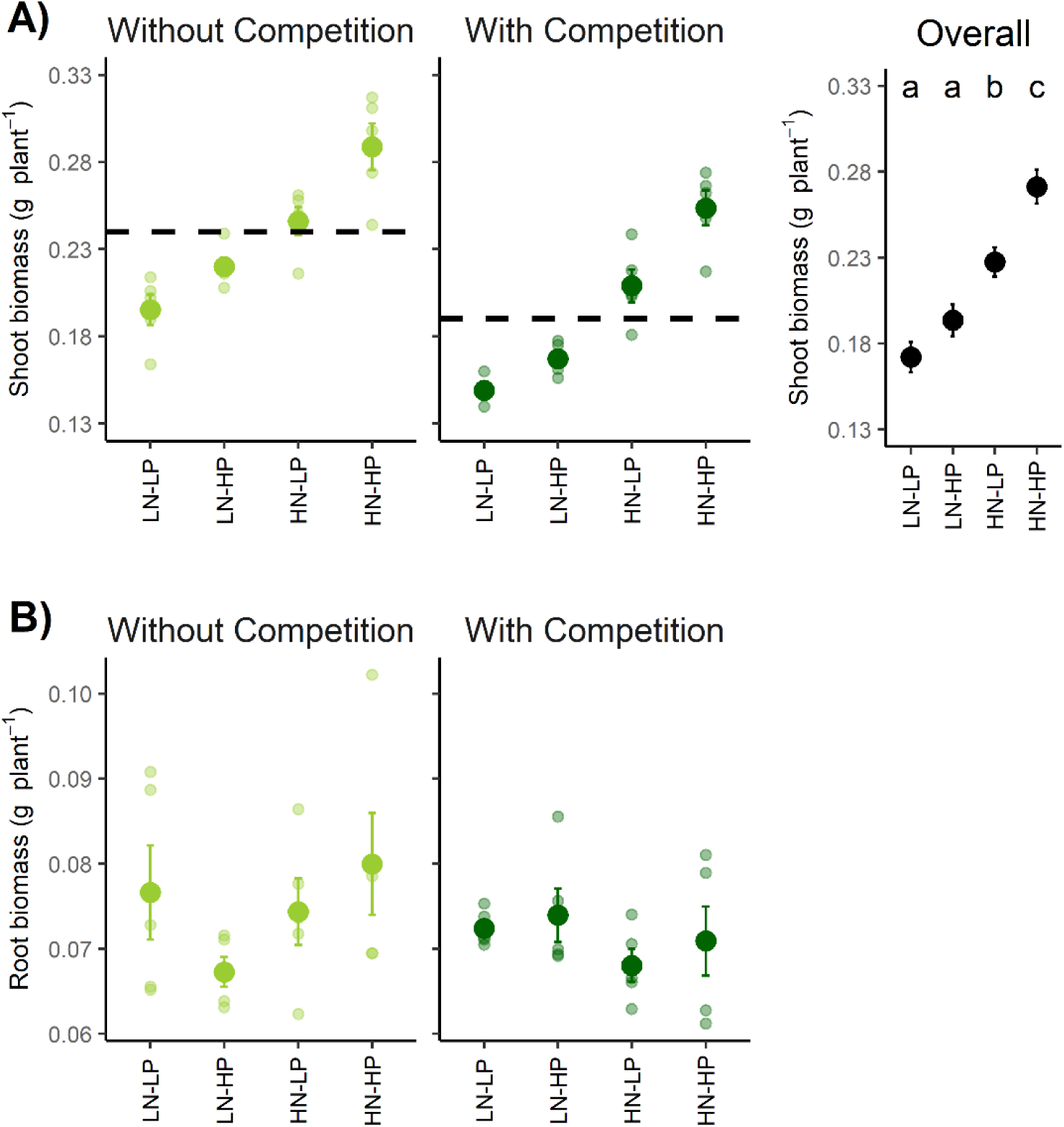
A) shoot and B) root biomass (g plant^-1^±SE) across N:P stoichiometry and with and without competition. LN-LP: low N and low P, LN-HP: low N and high P, HN-LP: high N and low P, and HN-HP: high N and high P. For shoot biomass, there was no interaction between N:P stoichiometry and competition. Therefore, a graph showing the results for each N:P stoichiometry level (across competition levels) is also displayed. For shoot biomass, dashed lines show mean shoot biomass values without and with competition. For each panel, different letters indicate significant differences (Tukey’s post-hoc, *P* < 0.05).

### Root system responses

Even though the greater amount of either N, P, or both increased shoot biomass, neither N:P stoichiometry (F_3,32_ = 0.79, *P* = 0.51) nor competition (F_1,32_ = 1.49, *P* = 0.24) had an effect on total root biomass production (Fig. 3B). However, biomass allocation as measured by the RMF was affected by N:P stoichiometry (F_3,32_ = 32.62, P < 0.001), competition (F_1,32_ = 26.01, P < 0.001), and their interaction (F_3,32_ = 5.77, P = 0.002). Irrespective of the presence or absence of competition, RMF was greater when both N and P were provided in low amounts (LN-LP) (Fig. 4). A high amount of either N, P, or both decreased RMF when plants were grown without competition. In contrast, when plants were grown in competition, providing high P (LN-HP) had no effect on RMF as compared to LN-LP (Fig. 4). In addition, providing high N with low or high P (HN-LP and HN-HP) reduced RMF both in the presence and absence of competitors.

**Fig. 4:**
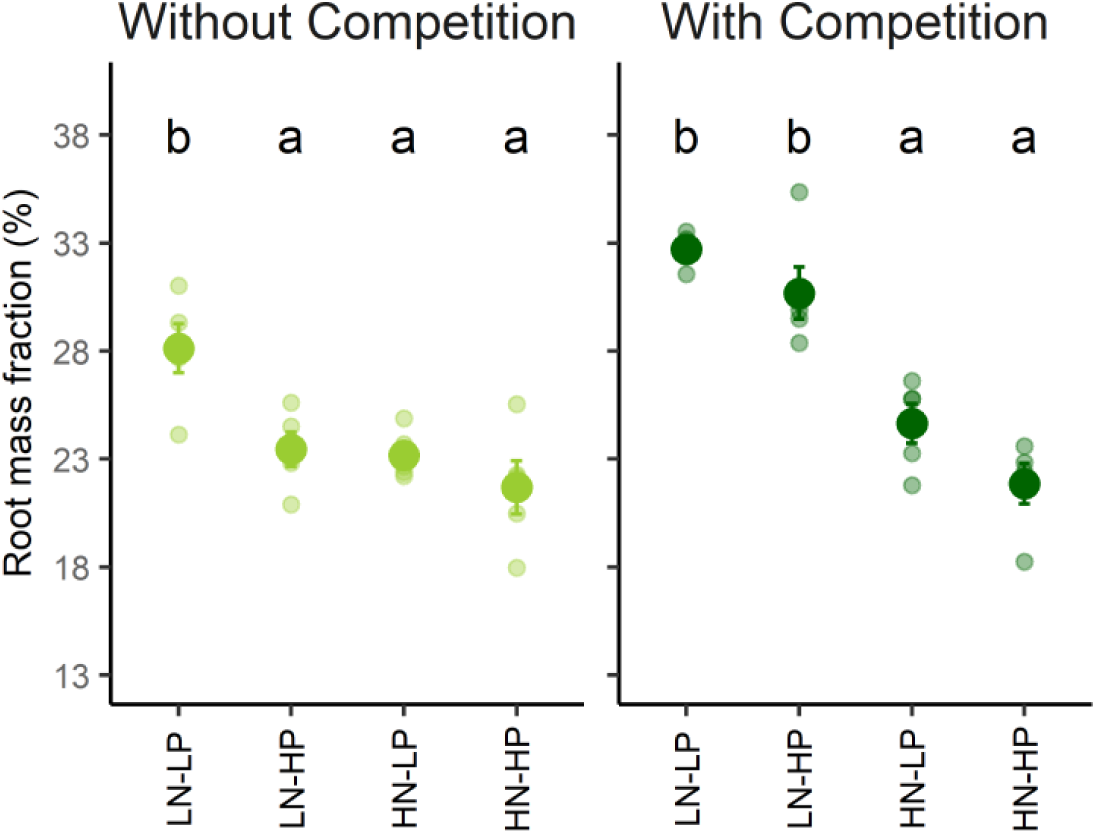
Root mass fraction (%) across N:P stoichiometry and with and without competition. LN-LP: low N and low P, LN-HP: low N and high P, HN-LP: high N and low P, and HN-HP: high N and high P. For each panel, different letters indicate significant differences (Tukey’s post-hoc, *P* < 0.05).

Vertical root distribution (*β*) was affected differently across N:P stoichiometry levels for plants growing alone or in competition (N:P stoichiometry: F_3,32_ = 22.19, P < 0.001; competition: F_1,32_ = 59.46, P < 0.001; N:P stoichiometry × competition: F_3,32_ = 4.85, P = 0.006). Vertical root distribution was governed by the identity of the nutrient being the most limiting (either N, P, or both) only for individually grown barley plants (in absence of competition). Without competition, plants rooted the deepest (greatest *β* value) when both N and P were provided in low amounts (LN-LP). On average, plants grown without competition in the LN-HP treatment also rooted deeper than the ones growing in the HN-LP and HN-HP treatments (Fig. 5A). Interestingly, the presence of competitors had a strong effect on the vertical root distribution measured at the population level. In this situation, the identity of the nutrient being the most limiting did not have any impact on root distribution. Overall, plants tended to increase their biomass allocation in roots to deeper soil layers (greater *β* values) when growing in competition (Fig. 5A).

**Fig. 5:**
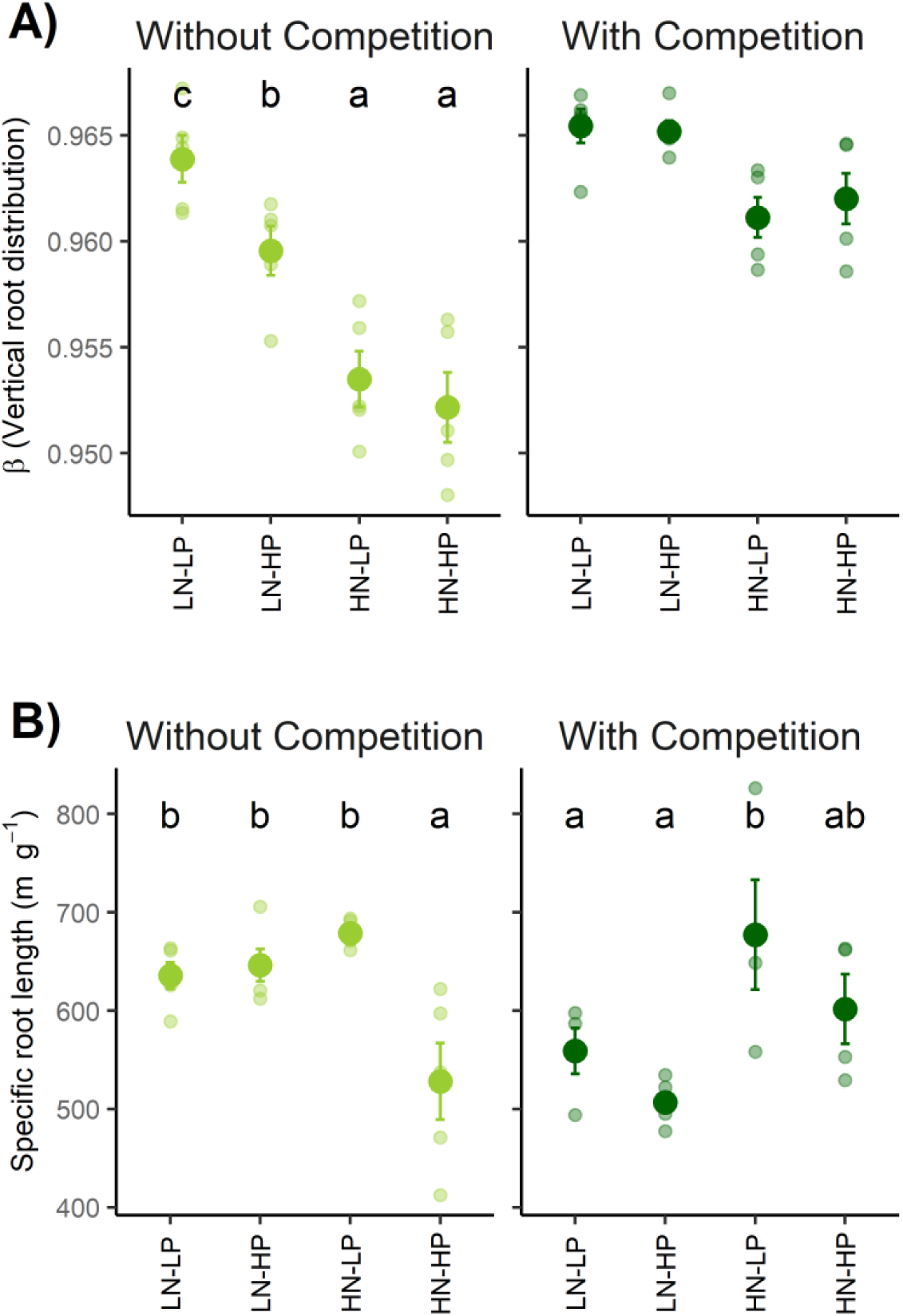
A) Vertical root distribution (β±SE, see methods) and B) specific root length (m g^-1^±SE) across N:P stoichiometry and with and without competition. LN-LP: low N and low P, LN-HP: low N and high P, HN-LP: high N and low P, and HN-HP: high N and high P. For each panel, different letters indicate significant differences (Tukey’s post-hoc, *P* < 0.05).

Even though the belowground biomass production remained similar between experimental treatments, root morphology was clearly impacted. Specific root length (SRL) was affected by N:P stoichiometry (F_3,32_ = 7.06, P = 0.001) and interacted with competition (F_3,32_ = 5.70, P = 0.003), but competition alone had no effect on SRL (Fig. 5B). Without competition, SRL was greater when either N, P, or both were provided in low amounts and did not depend on the identity of the nutrient being the most limiting. In contrast, in the presence of competition, SRL was greater only when P was the only limiting nutrient (HN-LP) (Fig. 5B).

### Shoot N:P concentrations

N:P stoichiometry and competition (presence/absence) had distinct effects on shoot N and P concentrations. Providing more N (HN-LP and HN-HP) or more P (LN-HP and HN-HP) resulted in greater shoot N and P concentrations, respectively. Shoot N concentration was significantly altered by N:P stoichiometry (F_3,32_ = 222.9, *P* < 0.001), competition (F_1,32_ = 259.3, *P* < 0.001), and their interaction (F_3,32_ = 10.9, *P* < 0.001) (Fig. 6A). Without competition, shoot N remained similar for both HN-LP and HN-HP, whereas, in the presence of competition, plant shoots had a greater N concentration under HN-LP than HN-HP (Fig. 6A). On the other hand, shoot P concentration was altered only by N:P stoichiometry (F_3,32_ = 9.19, *P* < 0.001). Providing more P increased its concentration in shoots on average by 53% (LN-HP) and 42% (HN-HP) (Fig. 6B). Our results also showed the existence of a positive correlation between SRL and shoot N concentration (no correlation between SRL and shoot P concentration), but only under intraspecific competition (Fig. 7). Further, shoot N:P mass ratio was affected by N:P stoichiometry (F_3,32_ = 21.72, P < 0.001) and competition (F_1,32_ = 5.50, P = 0.025). Compared to LN-LP, lower shoot N:P values were observed when N was the only limiting nutrient, while greater shoot N:P values were observed when P was the only limiting nutrient in the soil solution **[Supplementary Information Fig. S1]**. Shoot N:P ratios decreased from 24.6±2.3 to 20.1±2.1 in the presence of intraspecific competition.

**Fig. 6:**
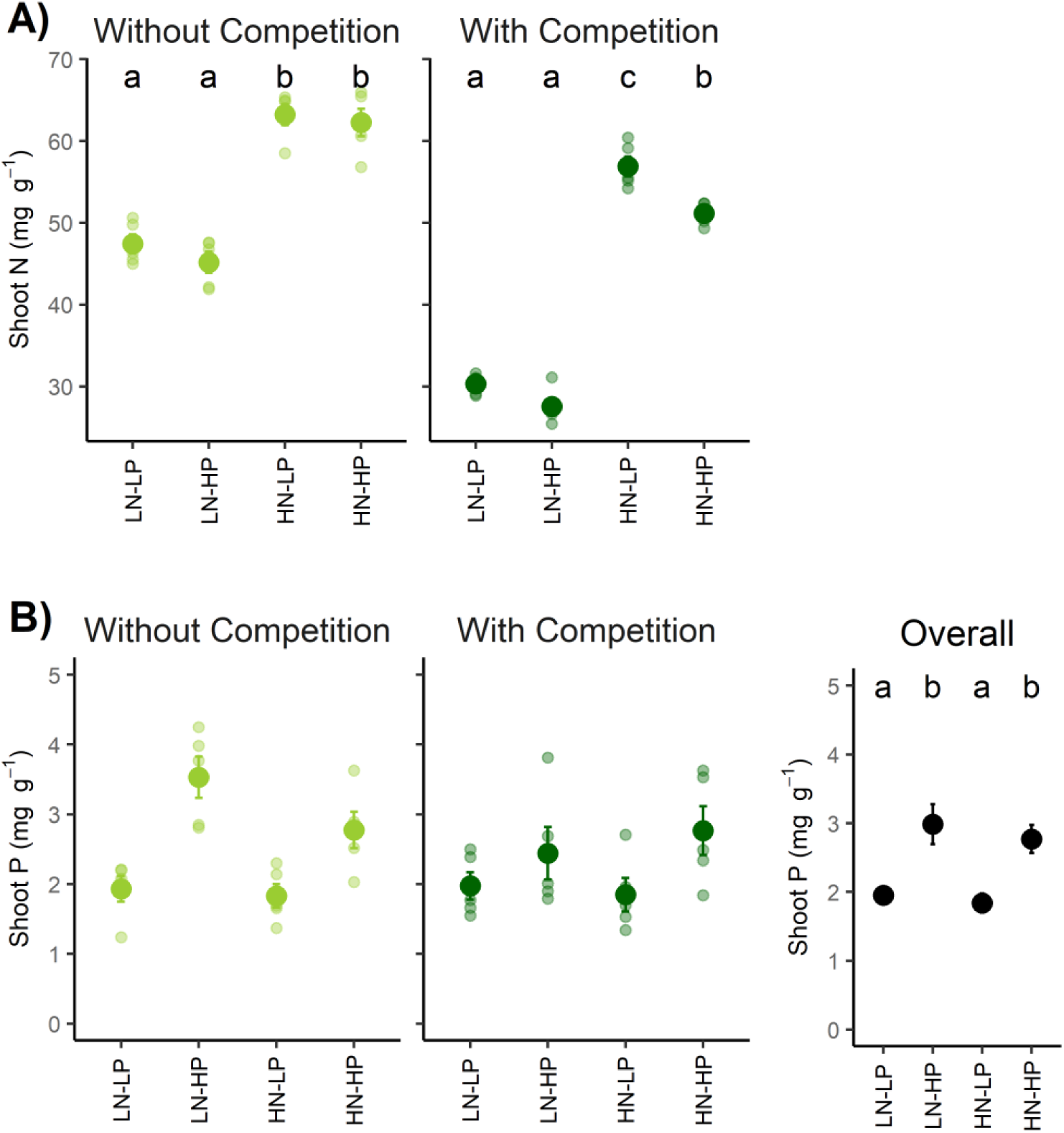
A) shoot N and B) shoot P (mg g^-1^±SE) across N:P stoichiometry and with and without competition. LN-LP: low N and low P, LN-HP: low N and high P, HN-LP: high N and low P, and HN-HP: high N and high P. For shoot P, there was no interaction between N:P stoichiometry and competition. Therefore, a graph showing the results for each N:P stoichiometry level (across competition levels) is also displayed. In each panel, different letters indicate significant differences (Tukey’s post-hoc, *P* < 0.05).

**Fig. 7:**
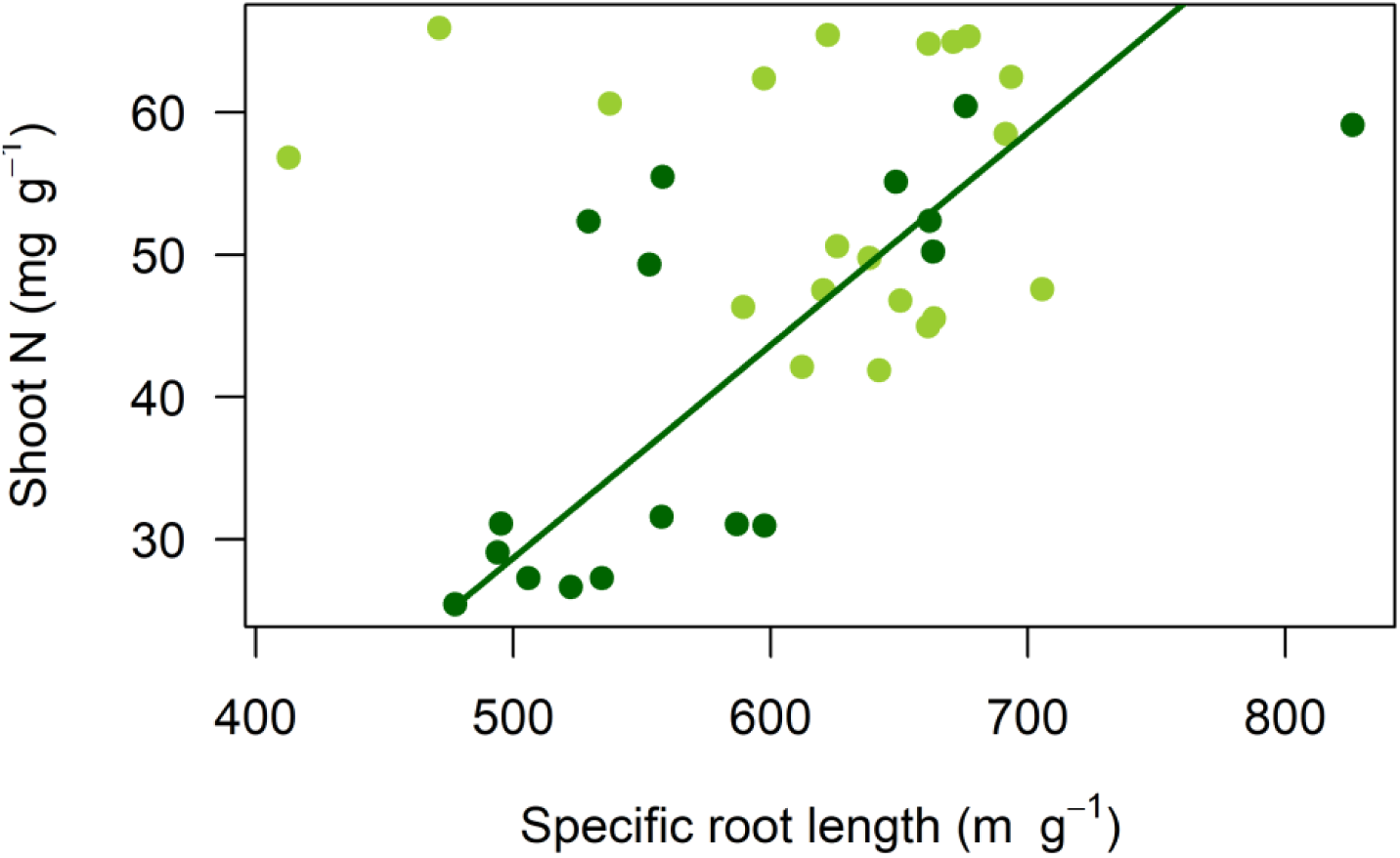
Linear relationship between specific root length (m g^-1^) and shoot N concentration (mg g^-1^). Light green circles represent absence whereas dark green circles represent presence of competition, respectively. Regression line between shoot N and specific root length is shown only when competition was present (for dark green).

## DISCUSSION

### Shoot but not root biomass production is more limited by N than by P

There is a general consensus that plants respond to nutrient shortage by changing their allocation patterns both below- and aboveground. When the availability of both macronutrients was low (LN-LP), aboveground productivity was the lowest, indicative of nutrient limitation. On the other hand, providing extra P or not, did not increase the shoot biomass production if N was the limiting nutrient (both in LN-LP and LN-HP), highlighting higher N demand for biomass production. Leaf N content is generally related to C assimilation during photosynthesis (Gastal and Lemaire, 2002). If reduced leaf N content leads to a reduction in the plant’s photosynthetic activity, a lower shoot biomass production can be expected when N is limiting in the soil (Fig.1). Andrew et al. (1999) showed for *Pisum sativum, Triticum aestivum*, and *Phaseolus vulgaris* that N shortage effects on plant growth are through its effects on protein synthesis. This further demonstrates that N limitation is more severe than P limitation for plant growth (see Capek et al. 2018) as the availability of extra P (LN-HP) in our study did not lead to higher shoot biomass production, probably due to N-mediated decrease in photosynthetic activity. Increased availability of both N and P (HN-HP), on the other hand, resulted in the greatest shoot biomass production because of greater N and P uptake that might ultimately lead to higher photosynthetic activity (Kumar et al. 2019).

Interestingly, root biomass production remained similar across N:P stoichiometry levels, but the RMF was greater when both N and P availability was low (LN-LP) in the absence of competition. This follows the general plant response to increasing C investment belowground when nutrient availability in the environment is low (Poorter et al. 2012).

Nutrient availability can strongly direct resource allocation patterns in plants (Gastal and Lemaire, 2002). More C allocation to roots under low nutrient availability is a well-known plant response as a potential mechanism to optimize growth by exploring a greater proportion of the soil volume for nutrients (de Groot et al. 2003; Hammond et al. 2006; Lambers et al. 2006). This is in line with optimal resource allocation theory, which predicts higher resource partitioning in organs that maximize the plant growth (Bloom et al. 1985). Increased RMF due to nutrient shortage allows plants to forage more effectively, yet it trades off with resource allocation in shoot biomass production (Garnett et al. 2009). We are aware that RMF only provides information about resource allocation to root growth component and does not necessarily include other carbon investments such as root respiration and exudation, yet it provides a hint about plant investments belowground for nutrient foraging. Greater availability of both N and P (HN-HP) has potentially led to lower investment belowground as shown in various studies for different vegetation (Aerts et al. 1991; Klimes and Klimesova, 1994; Wright et al. 2014). This further supports the notion of preferential uptake of available nutrients by roots, thereby minimizing their resource investments belowground for nutrient acquisition. These findings partly support our first hypothesis as the response to N limitation was only seen for the shoot but not root biomass.

### Intraspecific competition reduces shoot but not root biomass production

There is less debate with regard to the effect of competition (whether inter- or intraspecific) on biomass production, with several studies showing a decrease in plant biomass when growing in competition (Zhou et al. 2017; Heuermann et al. 2019). We also show that shoot biomass decreased in the presence of competition. A common underlying reason for this decline in biomass production when plants are competing is due to quick uptake of available nutrients leading to soil nutrient shortage (Tilman, 1990; Craine and Dybzinski, 2013). Surprisingly, we did not observe any change in root biomass production with or without competition. When plants are competing, and if plant growth is mostly affected by nutrient availability in soil, we would expect a greater resource investment in belowground organs to enhance nutrient uptake. In the presence of competition, a strong decrease in shoot biomass without altering root biomass per plant is confirmatory of increasing competitive ability for belowground resources, but at the expense of shoot biomass production. This also hints towards the plant’s phenotypic plasticity in biomass partitioning between shoots and roots. According to the competition model for limiting resources (Van Wijk et al. 2003), a lower investment belowground cannot sustain plant growth due to lower nutrient availability when plants are competing with each other. To maintain growth, therefore, higher investment in roots should be favored. In a recent study focusing on interspecific competition (growing oat with clover), increased root to shoot ratio without affecting shoot biomass production highlights that competition favored root biomass production for nutrient access (Heuermann et al. 2019). Further, the observed increase in RMF without affecting total root biomass under low N availability (LN-LP and LN-HP) supports our first hypothesis that N is more limiting plant growth than P limitation. Secondly, our second hypothesis is partly supported as only shoot biomass but not root biomass decreased with the intraspecific competition.

### Plants root deeper when limited by N, but only when growing without competitors

Root biomass may not always be indicative of the absorptive capacity of roots, and significant modifications in root morphology, anatomy, and architecture are possible with or without altering the total root biomass (Hodge, 2004). In our study, although the total root biomass remained similar between experimental treatments, we showed that the effect of N:P stoichiometry affected root system responses differently depending on the presence or absence of competitors. Such root system responses can be driven by relative mobility and, therefore, availability of N and P in soil strata. Vertical root distribution depended strongly on the identity of the limiting nutrient (either N, P, or both) in the absence of competition. For example, plants rooted shallower (lower *β* value) when P availability was low (HN-LP) whereas plants rooted deeper (higher *β* value) when N was the most limiting nutrient (LN-HP). Interestingly, when both nutrients were limiting (LN-LP), *β* was greatest thus suggesting that vertical root distribution was more likely driven by N limitation than P limitation and higher N than P demand. Given that P is less mobile than N in the soil matrix (Harrison, 1987), we expect more P to be present in the topsoil and more N to be present in deeper soil layers, and their relative limitations may have guided root responses. Plants respond to P shortage by reducing the primary root elongation but an increased number of lateral roots (Vance et al. 2003; Sanchez-Calderon et al. 2005). Further, Jia et al. (2018) showed that increasing the lateral root branching enhanced maize P acquisition. Gruber et al. (2013) also showed a shallower yet highly branched root system for *Arabidopsis* under P deficiency. On the other hand, when N is limiting plant growth, the plant’s investment in deep root systems is favored (Koevoets et al. 2016). In the presence of competition, β values did not change across N:P stoichiometry levels. Competition most likely resulted in quick nutrient uptake. Therefore, roots foraged throughout the rhizobox to their maximum extent to get excess to both N and P. In support of our third hypothesis, we show that plants root deeper when N is the most limiting nutrient, whereas shallower when P is the most limiting nutrient, but only in the absence of competition. Further, in the presence of intraspecific competition, root foraging is modulated by deeper soil exploration.

We also showed that, in the absence of competition, the SRL was greater when either N, P, or both were available in low amounts relative to HN-HP (Fig. 5B). Changes in SRL are general root morphological responses to lower availability of nutrients in the soil (Kong et al. 2014). By increasing SRL without altering the overall root biomass, plants are able to increase their foraging capacity. However, this may also be an apparent strategy of plants for nutrient acquisition as thinner roots have a lower life span and faster turnover (McCormack et al. 2012). On the contrary, when both N and P are not limiting plant growth (under HN-HP), it is more favorable for plants to invest less in increasing SRL due to associated aboveground allocation trades off. We expected the same effect of N:P stoichiometry on SRL in the presence of competition. However, we found contrasting effects, and SRL was lower when only N (LN-HP) or both N and P (LN-LP) were available in low amounts, whereas it increased only under HN-LP (high N but low P availability). As P is less mobile than N in the soil, increasing P foraging by greater SRL is likely one efficient strategy to increase its uptake. In contrast, greater N mobility would rather result in a deeper rooting system than increasing SRL locally to increase its uptake efficiently. Greater SRL with low P but high N availability (HN-LP) resulted in higher N uptake and associated higher P requirement. However, increased SRL did not result in higher P uptake due to its low availability. This further explains the positive relationship between SRL and shoot N uptake (probably as an indirect consequence of P limitation) (Fig. 7). These findings contrast strongly with results from a study in grasslands by Mommer et al. (2010), where interspecific competition with neighbors caused both higher investment of plants in root biomass as well as an accumulation of roots in the topsoil. This contrasting result could be driven by differences in root responses depending on whether neighbors are of the same or different species. Clearly, the presence of neighbors, whether of the same species or not, can drive this partly unexpected responses of roots. Whether experimental conditions are controlled (in the greenhouse) or not (in the field) will also probably affect the outcome.

### Effect of N:P stoichiometry and competition on shoot N and P concentrations

Shoot N and P concentrations were in line with what was expected. Providing high N (HN-LP and HN-HP) or high P (LN-HP and HN-HP) resulted in greater N and P concentrations in shoots, respectively. Intriguingly, in the presence of competition, we found that when both N and P availability was high (HN-HP), shoot N concentration was slightly lower than in plants grown under high N and low P (HN-LP) availability. This can most likely be explained by the fact that when both N and P were high, plants grew better (higher shoot biomass under HN-HP than HN-LP) and, as a consequence, exacerbated greater N demand. On the other hand, shoot P concentration was driven only by its availability in the soil and was similar for both with or without competition. This further supports our first hypothesis that soil N availability has a stronger effect in regulating plant performance more than P.

## CONCLUSIONS

Early plant responses to soil nutrient availability and plant-plant competition are decisive for plant performance. Lower shoot biomass under low N availability irrespective of P availability (both for LN-LP and LN-HP) indicates N limitation for shoot biomass production most likely due to higher N demand for photosynthesis. Higher investments belowground as a response to nutrient limitation pose a tradeoff with shoot biomass production. Roots foraged differently for N or P uptake by rooting deeper when N was limiting, but rooting shallower when P was limiting plant growth. However, when plants were competing for N and P in soil solution, no decrease in root biomass but lower shoot biomass per plant indicated differential resource allocation pattern by plants for maximizing nutrient uptake. When competing, plants rooted deeper indicating higher N demand and associated root acquisition strategy under these conditions. Such shift in plant resource allocation and root growth are key determinants for early plant nutrient acquisition and establishment, and illustrate the importance of biotic as well as abiotic drivers of plant responses to their environment. Field studies that manipulate N:P stoichiometry and focus on root foraging responses would move the field further forward now.

## ACKNOWLEDGMENTS

We thank Thomas Niemeyer for greenhouse assistance, Saatzucht Breun for supplying barley seeds free of charge, Dr. Kathleen Lemanski and Prof. Michael Bonkowski for arranging the Jackerath loess soil, Hannes Schempp, Hannah Uther, Johanna Wille, and IAESTE students for root scanning and laboratory assistance.

## DATA AVAILABILITY

Raw data and R scripts used for data analyses can be freely accessed at https://doi.org/10.5281/zenodo.3613623

---

**Supplementary figure S1:**
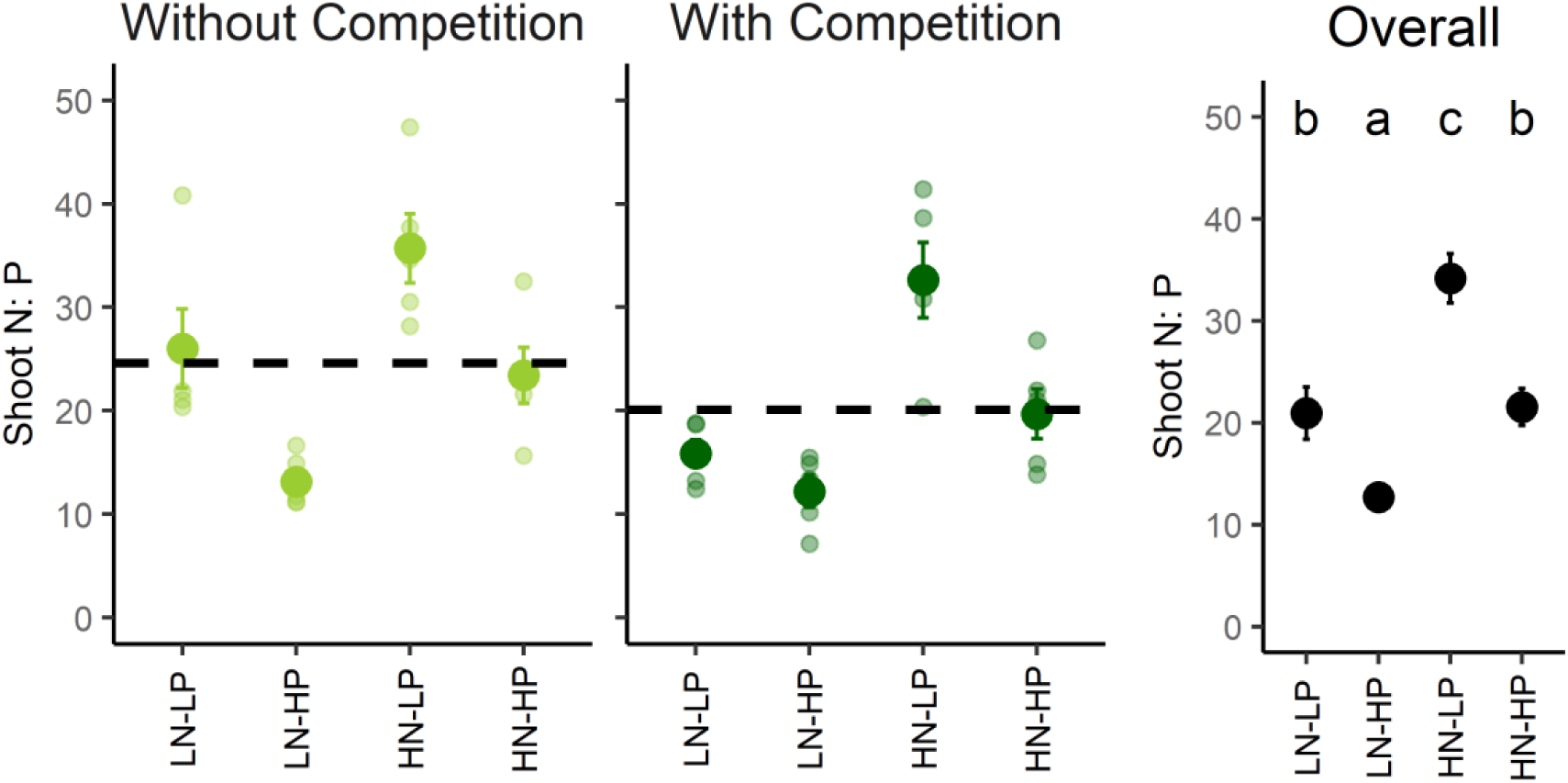
Shoot N:P mass ratio (±SE) across N:P stoichiometry and with and without competition. LN-LP: low N and low P, LN-HP: low N and high P, HN-LP: high N and low P, and HN-HP: high N and high P. For shoot P, there was no interaction between N:P stoichiometry and competition. Therefore, a graph showing the results for each N:P stoichiometry level (across competition levels) is also displayed. Dashed lines show mean shoot N:P values without and with competition. Different letters indicate significant differences (Tukey’s post-hoc, *P* < 0.05).

**Supplementary table 1:** Chemical concentration of nutrient solutions provided to plants. The N:P mass ratio of each stoichiometry of N and P is also provided in the last row of the table.

| **Macronutrients** | Stock (M) | Low N-Low P (LN-LP) | Low N-High P (LN-HP) | High N-Low P (HN-LP) | High N-High P (HN-HP) |
| --- | --- | --- | --- | --- | --- |
| KNO<sub>3</sub> | 1 | 0.625 | 0.625 | 2.875 | 2.5 |
| Ca(NO<sub>3</sub>)<sub>2</sub> · 4H<sub>2</sub>O | 1 | 0.625 | 0.625 | 2.125 | 2.5 |
| KH<sub>2</sub>PO<sub>4</sub> | 1 | 0.125 | 0.5 | 0.125 | 0.5 |
| MgSO<sub>4</sub> · 7H<sub>2</sub>O | 1 | 1.0 | 1.0 | 1.000 | 1.0 |
| **Micronutrients** | Stock (mM) |  |  |  |  |
| H<sub>3</sub>BO<sub>3</sub> · H<sub>2</sub>O | 46.3 | 0.5 | 0.5 | 0.5 | 0.5 |
| MnCl<sub>2</sub> · 4H<sub>2</sub>O | 9.2 |  |  |  |  |
| ZnSO<sub>4</sub> · 7H<sub>2</sub>O | 0.77 |  |  |  |  |
| CuSO<sub>4</sub> · 5H<sub>2</sub>O | 0.36 |  |  |  |  |
| MoO<sub>3</sub> (85% molybdic acid) | 0.01 |  |  |  |  |
| Fe-Na-EDTA | 50.12 | 0.5 | 0.5 | 0.5 | 0.5 |
| **Replacements** | Stock (M) |  |  |  |  |
| K<sub>2</sub>SO<sub>4</sub> | 0.5 | 2.25 | 1.875 |  |  |
| CaCl<sub>2</sub> * 2H<sub>2</sub>O | 1 | 1.875 | 1.875 |  |  |
| **N:P mass ratio** |  | **5.81** | **1.45** | **22.47** | **5.81** |

